# Protein contact map prediction using multiple sequence alignment dropout and consistency learning for sequences with fewer homologs

**DOI:** 10.1101/2021.05.12.443740

**Authors:** Xuyang Liu, Lei Jin, Shenghua Gao, Suwen Zhao

## Abstract

The prediction of protein contact map needs enough normalized number of effective sequence (Nf) in multiple sequence alignment (MSA). When Nf is small, the predicted contact maps are often not satisfactory. To solve this problem, we randomly selected a small part of sequence homologs for proteins with large Nf to generate MSAs with small Nf. From these MSAs, input features were generated and were passed through a consistency learning network, aiming to get the same results when using the features generated from the MSA with large Nf. The results showed that this method effectively improves the prediction accuracy of protein contact maps with small Nf.

## 1. Introduction

De novo protein structure prediction has been a long-standing challenge in computational biology. Since 2015, protein contact map assisted protein structure prediction has shown its great advantage and thus has been implemented in almost all cutting-edge *de novo* protein structure prediction tools, such as RaptorX^1^, I-TASSER^2^, AlphaFold1^3^, trRosetta^4^ and etc^5–8^.

Protein contact map is predicted by using features extracted from multiple sequences alignment (MSA). Such features include mutual information and direct-coupling analysis (DCA). Mutual information was often used in early stage studies to predict co-evolving residues pairs.^9–11^ However, mutual information includes both direct coupling pairs and indirect coupling pairs, while the latter are noise for protein contact prediction. To solve this problem, various DCA methods, such as mfDCA^12^, plmDCA^13^, EVfold^6,14^, PSICOV^15^, CCMpred^16^, and Gremlin^17^, were introduced about 10 years ago to denoise the indirect coupling pairs. Compared to mutual information, DCA greatly improves the accuracy of protein contact map prediction. However, both mutual information and DCA are highly rely on the number of effective sequences (Neff) in MSA. Generally, Neff = Sij(80%) is used; certain research group have own preferences, e.g. RaptorX used Neff (70%), Gremlin used sqrt(Neff)/L, Neff/L, and Neff/sqrt(L) [DeepMSA]. We chose to use Nf hereafter in this manuscript. However, both mutual information and DCA are highly rely on the number of non-redundant sequences in MSA. Each sequence is reweighted by 1divided by number of sequences > 80% identity (Sometimes, 70% sequence identity was also used). The sum of sequences’ weights divided by the sqrt of MSA’s length is usually used for measured the depth of MSA. The accuracy of protein contact map prediction increases with Nf. When Nf is smaller than 128, the contact map prediction becomes very challenging^18^.

Another strategy for protein contact map prediction is supervised machine learning, which uses 1D features (such as position specific scoring matrix (PSSM), secondary structure prediction, and relative solvent accessibility) and 2D features (such as DCA, mutual information pairwise potential, and covariance matrix) extracted from MSA as input of a neural network. In the early day, only shallow network architecture was used (such as MetaPSICOV^19^ and PconsC2^20^)_and they can outperform DCA methods. Later, much deeper network architectures such as ResNet^21^ are employed as they can capture higher-order residue correlation; and great breakthrough in contact map prediction accuracy was achieved by methods (such as RaptorX^7^) implemented such network. In these state-of-the-art machine learning methods, MSA plays an important role. High quality MSA helps to improve contact precision. For example, the involvement of metagenome data can help finding more homologous sequence from beyond the whole-genome.^22^ Methods taking advantage of metagenome data, such as TripletRes^23^ and MapPred^24^, show that a better MSA with enough homologs are useful for improving deep learning based contact prediction methods. Using different MSAs generated by different sequence databases and searching parameters can also help improve the predictions. RaptorX reported that the average 4 predictions according to 4 different MSAs is 1%~2% better than a single feature.^25^ Transform-restrained Rosetta reported that sometimes MSA is unnecessarily deep, they use a MSA subsampling and MSA selection methods that improves the precision by 1.5% and 2%~3% respectively. But for proteins with few homologs, the quality of predicted contact map is still quite challenging and needed to be improved.

Taking inspiration from the latest progress on the analysis of different object sizes in object detection, we propose a novel deep learning framework to handle this challenge. First, we make data augmentation for MSA with enough normalized number of effective sequence (Nf). The argumentation was done by randomly select part of MSA’s sequences that is proposed as MSA dropout, to do feature extraction. Then, features are learned from both original MSAs and MSAs dropout by a network branch called consistency learning that guide our network learning the difference between small Nf features and large Nf features. The results show our methods have much better contact map accuracy for proteins with small Nf, and at the same time, achieves state-of-the-art performance for proteins with large Nf.

## 2. Methods

### Definition for normalized number of effective sequence (Nf) in MSAs

Here we define the depth of MSA by calculating the normalized number of effective sequence (Nf):

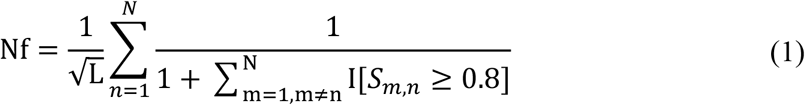

L is the number of residues in the protein sequence. N is the number of sequences in a MSA. *S_m,n_* is the sequence identity of *m*th sequence homolog and *n*th sequence homolog. I[*S_m,n_* ≥ 0.8] equals to 1 if *S_m,n_* ≥ 0.8, and to zero otherwise.

### Datasets selection

We made evaluation on two different test sets. The first test set is the same as described in Zhang’s article,^18^ which was from SCOPe database^26^ with 614 non-redundant protein. These 614 proteins have 403 easy and 211 hard targets, classified by the meta-threading program, LOMETS^27^. Here we mainly focused on the 211 hard targets. To further demonstrate the network’s performance on low Nf cases, we needed to test on more cases. So we used a subset of proteins from Protein Data Bank (PDB) with their first released date after December 2019. (Because the proteins released before December 2019 were used for training.) Using the PISCES website^28^, We removed the redundant proteins with a sequence identity larger than 25% to each other and resolution larger than 2.5 Å. Sequence was also ignored if its length was larger than 700 or less than 50. In this way, 1651 proteins from PDB were selected as the second test set.

Our training data was created from PDB in December 2019 using a subset of proteins which are satisfied the followings: (1) Sequence identity less than 25%. (2) Resolution less than 2.5 Å. (3) Sequence length is between 50 and 700. (4) Sequence identity large than 30% to any sequence in the 614 non-redundant proteins’ test sets were excluded.

### Multiple sequence alignment generation and sampling

The MSAs were generated using the Zhang lab’s DeepMSA software^18^. DeepMSA is a MSA generation pipeline by combining HH-suite^29^ and HMMER program^30^ to search homology, which can be divided into three stages in databases Uniclust, Uniref and Metaclust respectively. In this work, we generated MSAs using databases Uniclust30_2018_08^31^, Uniref90^32^ in December 2019 and Metaclust50_2018_08^33^. For each protein sequence in training and test datasets, the default search parameters in DeepMSA were used with the normalized number of effective sequence (Nf) cutoff 128. The Nf is non-redundant sequences with 80% sequence identity divided by the square root of the sequence length that is a commonly used approach in previous studies.

The sampled MSAs were also used for input feature generation. We randomly selected a part of homologs from the original MSA. The Nf interval of the sampled MSAs should be 10 to 20, ten MSAs were sampled. In a word, for each training protein sequence, 11 different MSAs were used (the original and 10 sampled MSAs) for input feature generation.

### Features generation

Input features are same with the RaptorX-Contact. Sequential and pairwise features were derived for every MSA. Sequential features (1D features) include protein position-specific scoring matrix (PSSM), predicted secondary structure, and solvent accessibility by RaptorX-property. Pairwise features include a DCA based contact map predicted by CCMpred, mutual information, and pairwise potential calculated by MetaPSICOV. For training proteins, both the original MSA and sampled MSAs were used for features generation. And for test proteins, only the original MSA was needed.

### Network architecture

The above process generated enough realistic small Nf cases from large Nf. However, simply learning from small Nf cases might not produce features discriminative enough. Taking inspiration from the recent work on contrastive learning networks^34^, we further proposed a feature-metric lifting loss to guide the training of the small Nf cases. In these frameworks, the authors proposed to learning a consistent feature representation from input data with different data augmentations to bootstrap the training of the network. Here, we followed a similar pipeline (**Figure 1**). Our intuition is that input features generated from a large Nf input are discriminative, and we want to learn a similar embedding from a small Nf input. Specifically, for each large Nf input *x_i_*, we first generated a small Nf pair 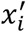 as in MSA sampling section, we then passed the input pair through a shared deep Resnet and enforced the learned consistency between these two inputs (Figure 1). We define the lifting loss as the l1 loss between the logits *z_i_* from *x_i_* and 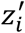 from 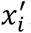.

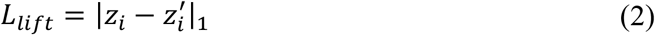

**Figure 1.**
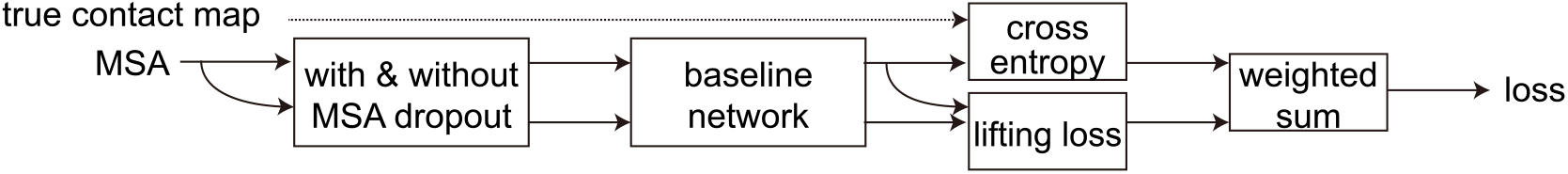
Architecture of our network.

On one hand, we proposed to learn a similar embedding between the small Nf inputs and large Nf inputs; on the other hand, the proposed lifting module further acts as a completion module, i.e., we supervised the network by completing a large Nf target with a small Nf input on the feature-metric level as in some recent works on image completion^34^. Note that our method is also a natural fit for unsupervised representation learning for protein contact map estimation. This will be our future research direction, i.e., to use the large number of unlabeled proteins for training deep conv-nets.

We reproduced RaptorX architecture according to Xu’s paper^1^. The 1D residual network consists of 30 1D residual blocks and 60 2D residual blocks. The convolution kernel size is 3 and we used different dilation to increase the receptive field for each neuron^35^. Batch normalization and ReLU activations were applied across different layers. Finally, a softmax layer was added to predicted the final output and adopt cross-entropy loss.

### Training

We used a two-step approach to test our proposal. We first pretrained the network on small Nf inputs only, then finetuned the network with the lifting loss. Our overall loss was a combination of the lifting loss and the standard cross entropy loss between our prediction and ground truth contact label. We implemented our solution under the PyTorch platform^36^. During training, we randomly sampled a 300*300 submatrix from the input sequence. The network was optimized end-to-end with AdamW optimizer^37^ for a total of 30 epochs with a batch size of 1. Learning rate was set to 1e-4 for the first 20 epoch. Then, we decayed it by 0.2 for every 5 epochs.

## RESULTS

### Dropout helps a lot for proteins of small Nf

Our model shows a great improvement on proteins of small Nf (5<Nf<40). We evaluated the performance of our model with TripletRes and RaptorX baselined on part of 211 hard targets cases that did not have enough homologous sequences. Here we show the results of proteins with effective sequence (Nf) between 5 and 40 with step of 5.

Generally, the contact precision is better with increasing Nf (**Figure 2a**). Although the performance of RaptorX baseline model is poor than TripletRes, the model trained by MSAs dropout and consistency learning has a significant improvement and is better than TripletRes on most cases. In summary, for targets with Nf between 5 and 40, network trained by the original MSAs performs not very well on the top L/5 precision. Using the same data, our model using consistency learning and MSAs dropout can improve the precision of proteins of small Nf from 0.727 to 0.818, and it’s better than TripletRes’ 0.779 (**Table S1**). The precision matrix, used by TripletRes, showed ability to perform better for proteins with low number of homologous sequences^23,38,39^, which was not used by RaptorX. This may lead to the poor performance of our RaptorX baseline. Nevertheless, used by consistency learning and MSA dropout, network can have better performance on these targets.

**Figure 2.**
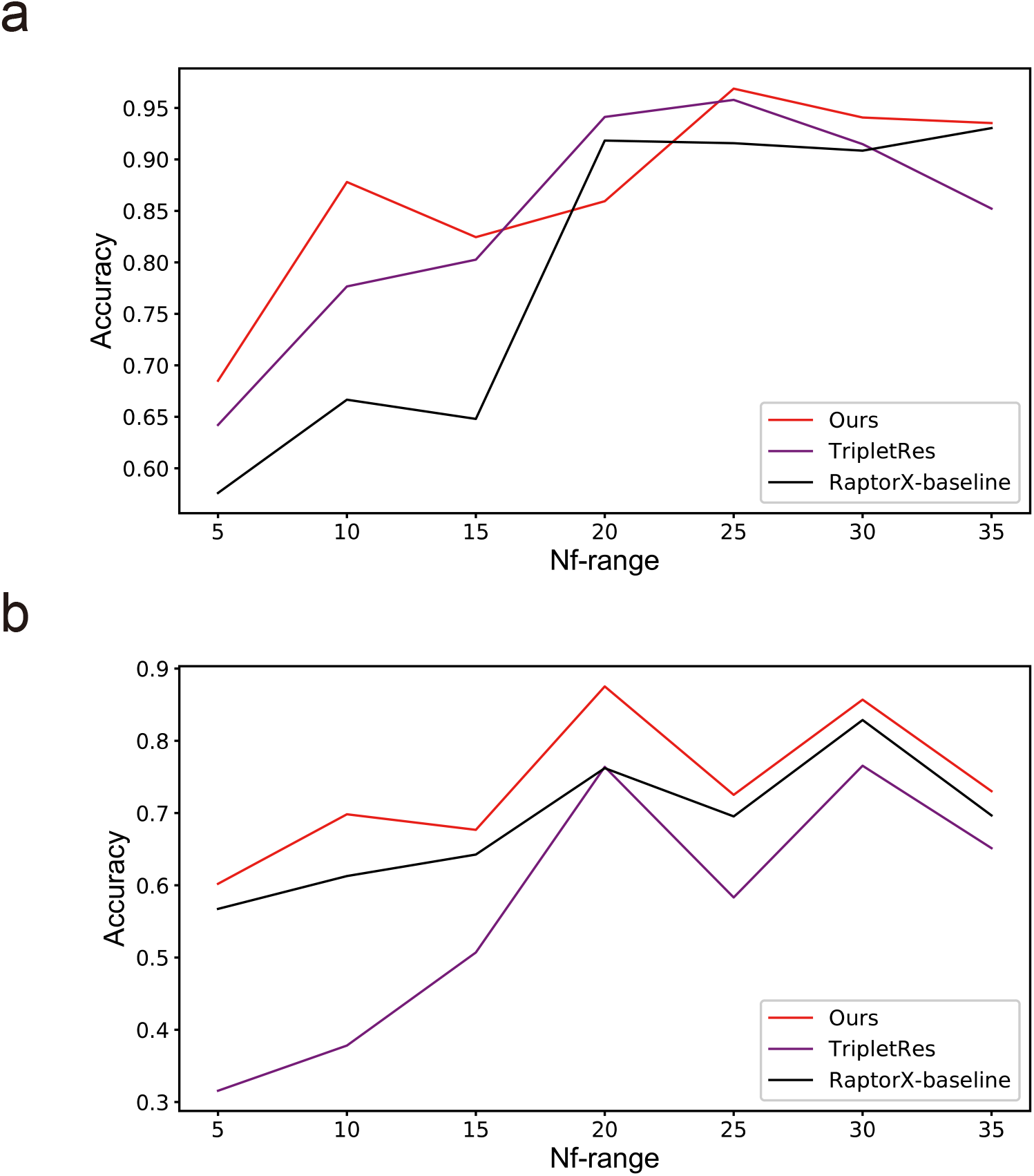
Performance of top L/5 long range contact on proteins with small Nf (5 <= Nf <= 40).

One thing needs to mention is the above TripletRes results was derived from Zhang lab’s DeepMSA article in 2020. The final performance can be influenced by many reasons such as the quality of multiple sequence alignments. We also want to know the performance of our model, TripletRes, and RaptorX baseline when using the same alignments. So, we make another test set consist of proteins that their structures are released later than December 2019. 1651 proteins were filtered as described in the methods. TripletRes results of these proteins are evaluated using standalone package of TripletRes with the same MSA as our model. We show the comparison of our model, TripletRes, and RaptorX baseline on small Nf cases. (**Figure 2b**). For almost all intervals, our trained RaptorX baseline performs better than TripletRes, while the model with data argumentation has even better performance. Just using the MSAs dropout and consistency learning network, the overall predicted precision of these proteins with Nf ranging from 5 to 40 improves from RaptorX baseline’s 0.656 to 0.707.

### Performance on large Nf proteins is comparable

Our methods achieve not only better performance on protein with small Nf, but also stat-of-art performance on large Nf proteins. Here we show the contact precision with enough homologs (Nf > 40) compared with TripletRes (**Table 2**). For 1651 test PDBs, the top L/5 long range precision of TripletRes, Reproduced RaptorX baseline and our methods are 0.918, 0.922, and 0.905 respectively. These models’ performances on large Nf are very close, and much higher than proteins with smaller Nf. On these proteins, our reproduced RaptorX baseline is comparable to the TripletRes, while the network with MSAs dropout has slightly lower performance that may be caused by balancing the information between dropped MSA and original MSA, suggesting that we should train different models for different Nf’s proteins to get the best performance.

**Table 1.** Small Nf performance of PDB test set.

| Nf-range | counts | ours | TripletRes | RaptorX baseline |
| --- | --- | --- | --- | --- |
| 5-10 | 42 | **0.602** | 0.316 | 0.567 |
| 10-15 | 24 | **0.698** | 0.378 | 0.613 |
| 15-20 | 22 | **0.677** | 0.507 | 0.643 |
| 20-25 | 15 | **0.875** | 0.764 | 0.762 |
| 25-30 | 11 | **0.725** | 0.583 | 0.695 |
| 30-35 | 15 | **0.857** | 0.765 | 0.829 |
| 35-40 | 12 | **0.730** | 0.651 | 0.697 |
| sum | 141 | **0.707** | 0.501 | 0.656 |

**Table 2.**
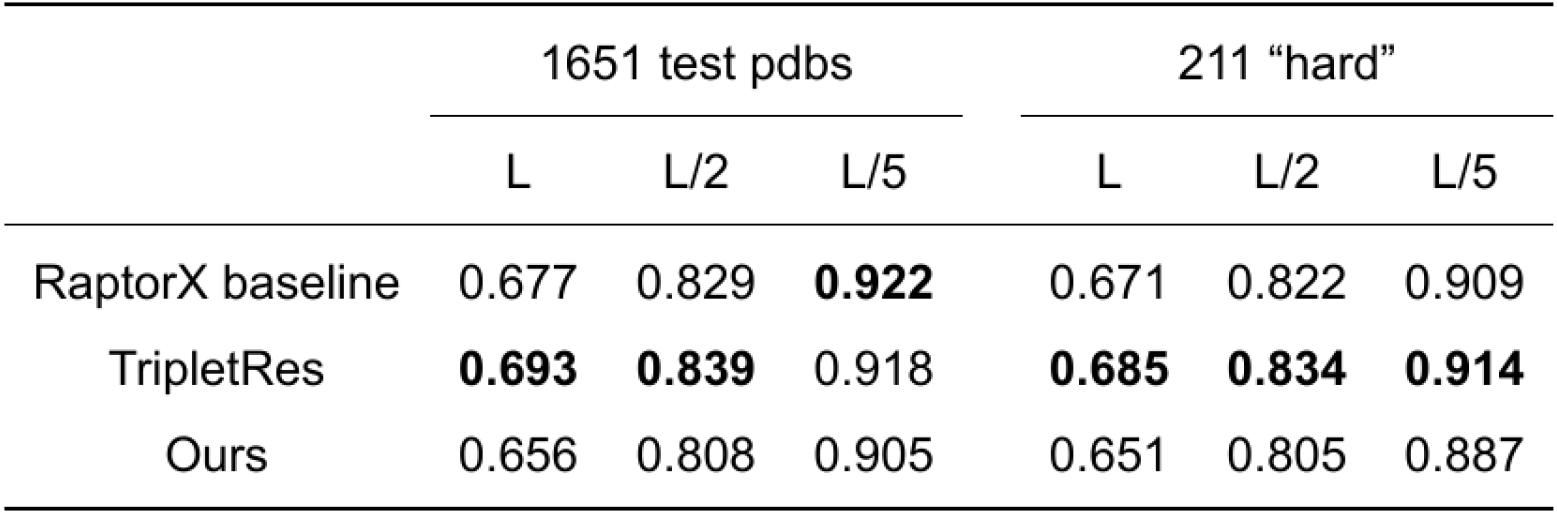
Long range precision for proteins with Nf > 40.

|  | 1651 test pdbs |  |  | 211 "hard" |  |  |
| --- | --- | --- | --- | --- | --- | --- |
|  | L | L/2 | L/5 | L | L/2 | L/5 |
| RaptorX baseline | 0.677 | 0.829 | **0.922** | 0.671 | 0.822 | 0.909 |
| TripletRes | **0.693** | **0.839** | 0.918 | **0.685** | **0.834** | **0.914** |
| Ours | 0.656 | 0.808 | 0.905 | 0.651 | 0.805 | 0.887 |

### Ablation Experiments

Here, we provide an analysis with each of the proposed module. We divided our test sets into three different subsets: Nf < 10, 10 <= Nf < 40, and Nf >= 40.

First, we analyzed the results of applying MSAs dropout module to the network. We report the results with long range top L/5 precision (**Table 3**). The first row shows the result when the original MSA and the original framework of RaptorX are used for training. The second row shows the performance when only MSA dropout is used for data argumentation. Using of MSA dropout, that is only 10 sets of small Nf data, increase 2~3% for data in the interval of Nf < 10 and 10 <= Nf < 40. We believe this is due to the obvious difference between small Nf data and large Nf data. In other words, small Nf data has fewer effective sequence homologs, so its input features do not have enough information. The previous studies ignored the difference between different Nf data. When small Nf data and large Nf data are trained simultaneously in the same network framework, the network will be more inclined to fit one type of data. In the current situation, the large Nf data in the training set accounts for the majority, so it is more inclined to fit the large Nf data, thereby reducing the accuracy of the small Nf data. So when we use more small Nf data to train the network, the network can have a better fit to the small Nf data..

**Table 3.** Ablation experiment.

| Accuracy(L/5) | 0< Testset Nf <=10 | 10< Testset Nf <=40 | Testset Nf > 40 |
| --- | --- | --- | --- |
| RaptorX Baseline | 0.471 | 0.694 | **0.922** |
| MSAs dropout | 0.496 | 0.721 | 0.893 |
| MSAs dropout + consistency learning | **0.533** | **0.751** | 0.905 |

The third row shows the performance when consistency learning network is added. For small Nf data, the performance is about 5% higher than the baseline network. This also illustrates the idea of using large Nf data to “guide” small Nf data, and the network has indeed learned the difference between these two types of data. Thus described, MSA dropout and consistency learning can be used to enhance small Nf protein’s contact map prediction. After adding the training process of MSA dropout and the consistency learning network, the precision is slightly lower by 1~2%. We believe this is due to the need to balance the features of small Nf inputs and large Nf inputs during the network training process, ignoring the learning weight of some high-Nf input features.

### Case study

To further compare the difference between our method and the baseline, we show several representative examples of small Nf. The visual representation of its contact map and three-dimensional conformation show the reliability of our method (**Figure 3**).

**Figure 3.**
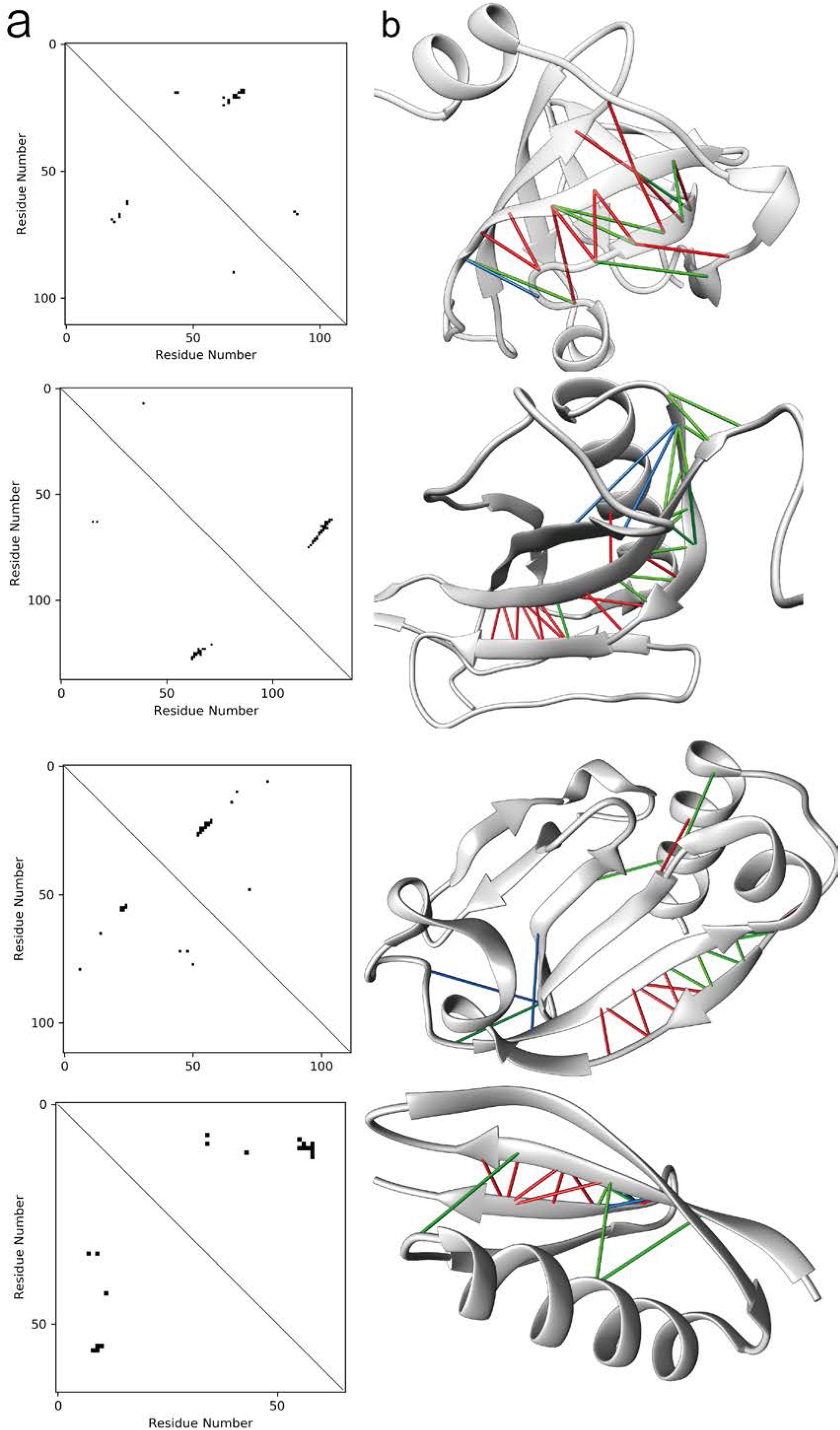
Case study of truly predicted contact of long L/5. (**a, b**) Contact map and 3-D structure of proteins d4l5qa1, d1qqra_, 6LYJC and 6OC7C from top to bottom. The lower triangle is the truly predicted contacts by baseline and the upper triangle is the truly predicted contacts with argumentation (**a**). Green lines are the top L/5 contacts predicted both by baseline and argumentation. Blue lines are contact only predicted by baseline and red lines are contacts only predicted by argumentation (**b**).

In general, the truly predicted top L/5 long range contacts by MSAs dropout and consistency learning cover most predicted residues by the original network. The first two examples are from SCOPe test set, the top L/5 long range contact of d4l5qa1 improves from 0.318 to 0.818, and d1qqra_ improves from 0.666 to 1.000. The last two examples are from PDB test set, 6LYJC improves from 0.500 to 0.818 and 6OC7C improves from 0.538 to 0.923. The argumentation network not only covers more regions of query proteins, but also finds more contacts around predictions of original network and reduced the false positive rates.

### Summary of strategies to improve protein contact map prediction by squeezing MSAs

We summarized some studies that improved the accuracy of contact map prediction by squeezing information from MSAs. These studies can be roughly divided into two categories. One is to increase the effective number homologs, the other is to subsampling sequences from MSAs (**Table 4**).

**Table 4.** Squeezing information in MSAs.

| Methods | Action | Improvement | References |
| --- | --- | --- | --- |
| RaptorX | Varying databases and parameters | 1%~2% | Wang *et al.* 2016 |
| trRosetta | MSA subsampling and selection | 3.5%~5% | Yang *et al.* 2020 |
| DeepMSA | Metagenome data | 2%~6% | Zhang *et al.* 2020 |
| Ours | MSA dropout and consistency learning | 5%~8% |  |

In 2016, RaptorX improved the prediction accuracy by using different sequence databases (UniProt20_2016_02 and UniProt20_2015_11) and different search parameters (E-value 0.001 and 1). Each sequence produced 4 input features by different MSAs^25^. This approach is equivalent to increase the Nf of MSAs and subsampling them, so it can improve the prediction accuracy of 1~2%. In CASP13, TripletRes, using metagenome sequence database to find more sequence homologs, enriched the information contained in MSA. This improves the prediction performance of small Nf^18,23^.

However, yang et al found that the more effective sequence homologs is not always better, but the reliability of sequence homologs need to ensure. They performed selection and subsampling for MSA. The selection of MSA refers to selection homologous sequences with low E-value and high coverage as much as possible, while the subsampling of MSA refers to only extracting 50% sequences from the selected MSAs in every training epoch. Through the combination of these two operations, the prediction performance improved 2~5%.

Here we mainly focus the small Nf proteins. We believe that one reason for its poor prediction effect is that there are fewer small Nf proteins in the training data. Mixing small Nf and large Nf input features makes the network pay more attention to large Nf features. Therefore, we artificially generate small Nf input features so that the network can learn the features of this part of the protein much better. In addition, we also found that large Nf protein can guide the prediction of small Nf protein. Finally, we increased the performance by 5% and 8% in two different test sets respectively.

## Conclusion and discussion

We proposed a novel data argumentation method for feature generation called MSAs dropout and implement consistency learning network in contact map prediction. We reproduced RaptorX contact prediction architecture and used MSAs dropout and consistency learning on it. We evaluate the performance with TripletRes, one of the best methods in contact map prediction in CASP13. Even the precision matrix used by TripletRes can greatly improve the precision of low Nf, our method, only use the PSSM and protein 1-D property instead, outperforms TripletRes. Meanwhile, our methods can achieve stat-of-art performance on protein with large Nf. So, we prove that MSAs dropout and consistency learning network is useful for contact map prediction.

Although we are using RaptorX’s contact map prediction architecture in this study, MSAs dropout and consistency learning network are general methods that can be combined with distinct MSA generated features and network architectures in protein contact/distance predictions and other protein property prediction.

In this study, we just used a range of MSAs dropout 10 ~ 20 that results in significant improvement for protein with relatively low Nf. For proteins with very small Nf, such as Nf less than 1, the features now contain little information for networks to infer the contact, more works will be done in future work.

---

**Figure S1.**
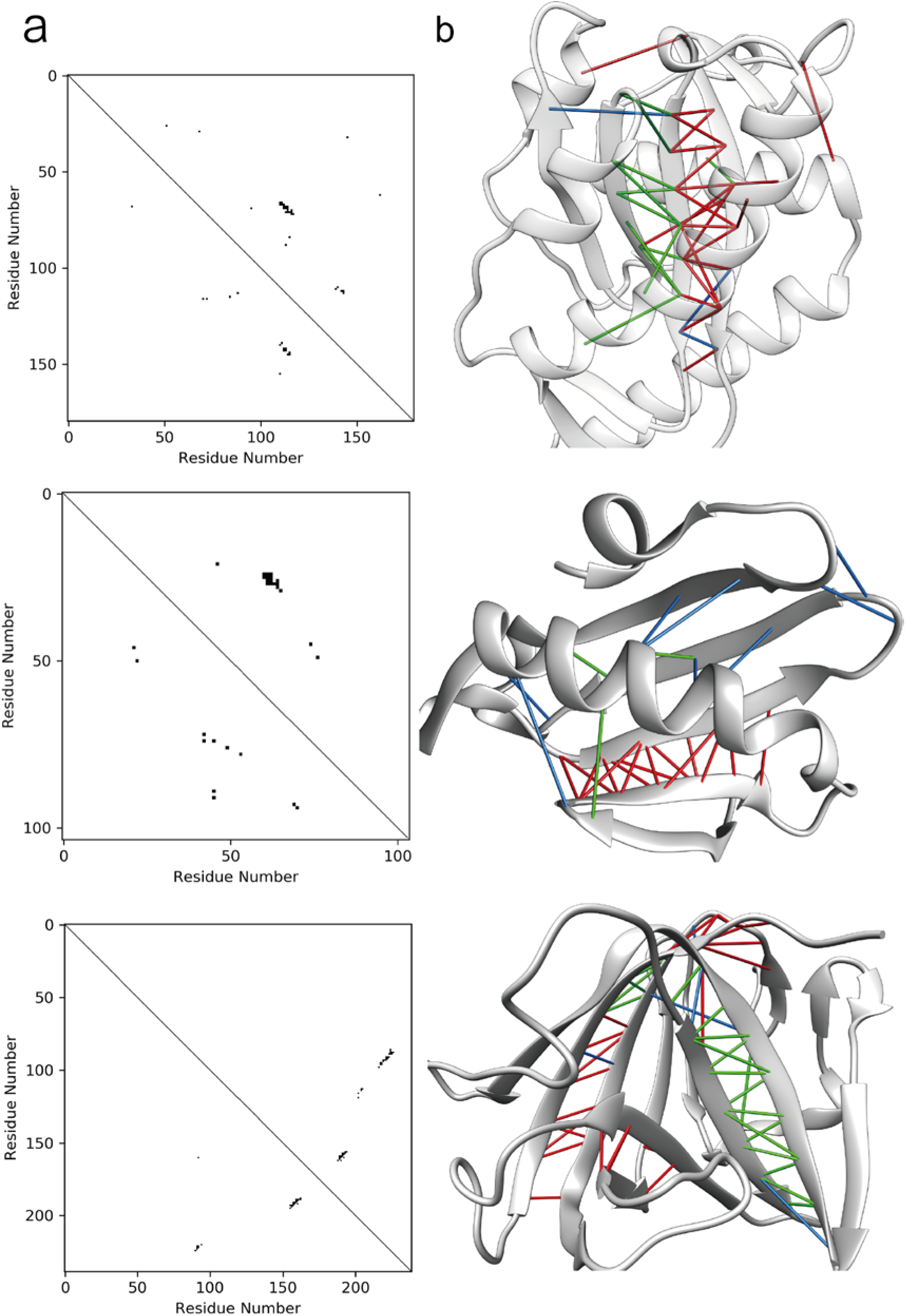
Case study of truly predicted contact of long L/5. (**a, b**) Contact map and 3-D structure of proteins d1y0ka1, 6UTCA and 7BV0A, from top to bottom. The lower triangle is the truly predicted contacts by baseline and the upper triangle is the truly predicted contacts with argumentation (**a**). Green lines are the top L/5 contacts predicted both by baseline and argumentation. Blue lines are contact only predicted by baseline and red lines are contacts only predicted by argumentation (**b**).

**Table S1.** Small Nf performance of scope test set.

| Nf-range | counts | ours | TripletRes | RaptorX baseline |
| --- | --- | --- | --- | --- |
| 5-10 | 15 | **0.685** | 0.642 | 0.576 |
| 10-15 | 5 | **0.878** | 0.777 | 0.667 |
| 15-20 | 7 | **0.825** | 0.803 | 0.648 |
| 20-25 | 4 | 0.859 | **0.941** | 0.918 |
| 25-30 | 2 | **0.969** | 0.958 | 0.916 |
| 30-35 | 3 | **0.941** | 0.915 | 0.909 |
| 35-40 | 7 | **0.935** | 0.852 | 0.931 |
| sum | 43 | **0.801** | 0.779 | 0.726 |

